# Interrogation of the glycolytic axis in pulmonary arterial hypertension models and how modifying these can lead to potential therapeutic options

**DOI:** 10.64898/2026.01.14.699109

**Authors:** Rachel Sutcliffe, Iona Cuthbertson, Stephen Moore, Mark Southwood, Benjamin Dunmore, Paul Upton, Nicholas Morrell, Paola Caruso, Elaine Soon

## Abstract

**Introduction:** Pulmonary arterial hypertension (PAH) describes diseases characterized by increased pulmonary vessel pressures that lead to right ventricular failure and death if left untreated. Dysregulated metabolic function has been reported in endothelial cells from PAH patients. Healthy endothelial cells can rapidly shift from quiescence to proliferative and apoptotic states, partly due to a greater utilization of glycolysis for ATP production, rather than the slower oxidative phosphorylation more commonly relied on by other cell types. However, in PAH patients, pulmonary artery endothelial cells demonstrate an even greater shift towards glycolytic ATP production. This phenomenon, which was first described in cancer as the ‘Warburg effect’, is an adaptation for optimizing both energy production and biosynthetic processes for maximum proliferative potential.

**Hypothesis and Methods:** We hypothesize that targeting the PKM2 axis in endothelial cells may shift the balance to favor oxidative phosphorylation, in effect ‘reversing’ the Warburg effect; and that this would have beneficial effects in cell and rodent models of PAH. To this end we have tested TEPP-46, an allosteric PKM2 activator that stabilizes the tetrameric form, blocking its translocation into the nucleus and increasing canonical enzymatic activity, in blood outgrowth endothelial cells (BOECs) and in a rat Sugen-hypoxia model.

**Results:** TEPP-46 treatment reduced levels of nuclear PKM2 (the homodimer form which promotes anabolic effects and proliferation). Cytoplasmic levels of PKM2 and levels of PKM1 were unaffected. Exposure to TEPP-46 lowered levels of polypyrimidine tract-binding protein 1 (PTPB1), which controls alternative splicing of the *PKM* gene to promote PKM2 expression; and reduced LDHA, which converts pyruvate to lactate, preventing its utilization for oxidative phosphorylation. In the Sugen-hypoxia rat model, administration of TEPP-46 significantly ameliorated the elevated right ventricular systolic pressures, reduced the loss of body weight and increased survival. The increased muscularization of the smaller blood arteries and arterioles in the lungs of rats due to Sugen-hypoxia were also improved by concurrent administration of TEPP-46.

**Conclusion:** We have shown that TEPP-46 treatment reduces nuclear PKM2 and represses PKM2-driven gene expression, with the functional effect of ameliorating the Sugen-hypoxia phenotype. This suggests that antagonizing the Warburg effect may offer a novel therapeutic avenue for PAH.

---

Pulmonary arterial hypertension (PAH) describes diseases characterized by high pulmonary pressures (greater than 20mmHg)(1) that lead to right ventricular failure and eventually death. This is due to remodeling of the lumens of small-to-medium pulmonary vessels, with eventual obliteration. PAH development is multifactorial, with multiple ‘hits’ required for initiation and pathogenesis, akin to carcinogenesis. Mutations in *BMPR2*, pro-inflammatory states, and endothelial cell metabolic abnormalities(2) have all been implicated as contributory factors.

Healthy endothelial cells are able to rapidly shift from quiescence to proliferative and apoptotic states, partly due to a greater utilization of glycolysis for ATP production, rather than the slower oxidative phosphorylation more commonly relied on by other cell types (3). However, in PAH patients, pulmonary artery endothelial cells demonstrate an even greater shift towards glycolytic ATP production(4). This phenomenon, which was first described in cancer as the ‘Warburg effect’, is considered to be an adaptation for optimizing both energy production and biosynthetic processes for maximum proliferative potential(5).

Pyruvate kinase plays a critical role in glycolysis, as this enzyme catalyzes phosphoenolpyruvate to pyruvate, which then either fuels oxidative phosphorylation, through the citric acid cycle, or is converted to lactate. There are four isoforms of pyruvate kinase, with the crucial two in endothelial cells being PKM1 and PKM2. These two isoforms arise from alternative splicing of the *PKM* transcript – inclusion of exon 9 results in PKM1 and inclusion of exon 10 results in PKM2. PKM2 is mainly expressed in embryonic cells and is progressively replaced by PKM1 in adult tissue, especially in cells requiring rapid energy provision, such as neurons and muscle. However, in tumors this reverts, with PKM2 again becoming the predominant isoform.

PKM2 shifts between multiple forms with different functions; a highly enzymatically active tetrameric form, which converts phosphoenolpyruvate into pyruvate, and a less active dimeric form, which can translocate to the nucleus and regulate the expression of numerous proteins (Fig.1A)(6). Due to the limited enzymatic activity of dimeric PKM2, this isoform, when prevalent, slows the production of pyruvate, leading to build-up of glycolytic intermediates that are re-directed into anabolic pathways.

We hypothesize that targeting the PKM2 axis in endothelial cells may shift the balance to favor oxidative phosphorylation, in effect ‘reversing’ the Warburg effect; and that this would have beneficial effects in cell and rodent models of PAH. To this end we have tested TEPP-46, an allosteric PKM2 activator that stabilizes the tetrameric form, blocking its translocation into the nucleus and increasing canonical enzymatic activity(7).

Blood outgrowth endothelial cells (BOECs) were selected as the cell model as these share similar characteristics to mature pulmonary artery endothelial cells, with only 0.005% of genes differentially expressed between them(8). BOECs were treated with either vehicle or TEPP-46 for 48 hours. After exposure to TEPP-46, PKM2 levels in the nuclear fraction (where the proliferation-promoting homodimeric form is present) was decreased (Fig.1B-C) while levels in the cytoplasm (where both forms exist) remain unchanged (Fig.1D-E). To explore the mechanism behind this, whole cell lysates were also immunoblotted. TEPP-46 treatment reduced levels of Polypyrimidine tract-binding protein 1 (PTPB1), which controls alternative splicing of the PKM gene to promote PKM2 expression (Fig1.F-G); and lowered levels of LDHA, which converts pyruvate to lactate, preventing its utilization for oxidative phosphorylation (Fig1.H-I). Levels of PKM1 were unchanged after TEPP-46 treatment (Fig.1J-K); which shows that the TEPP-46 treatment specifically affects PKM2. This implies that off-target effects of TEPP-46 are minimal.

**Figure 1:**
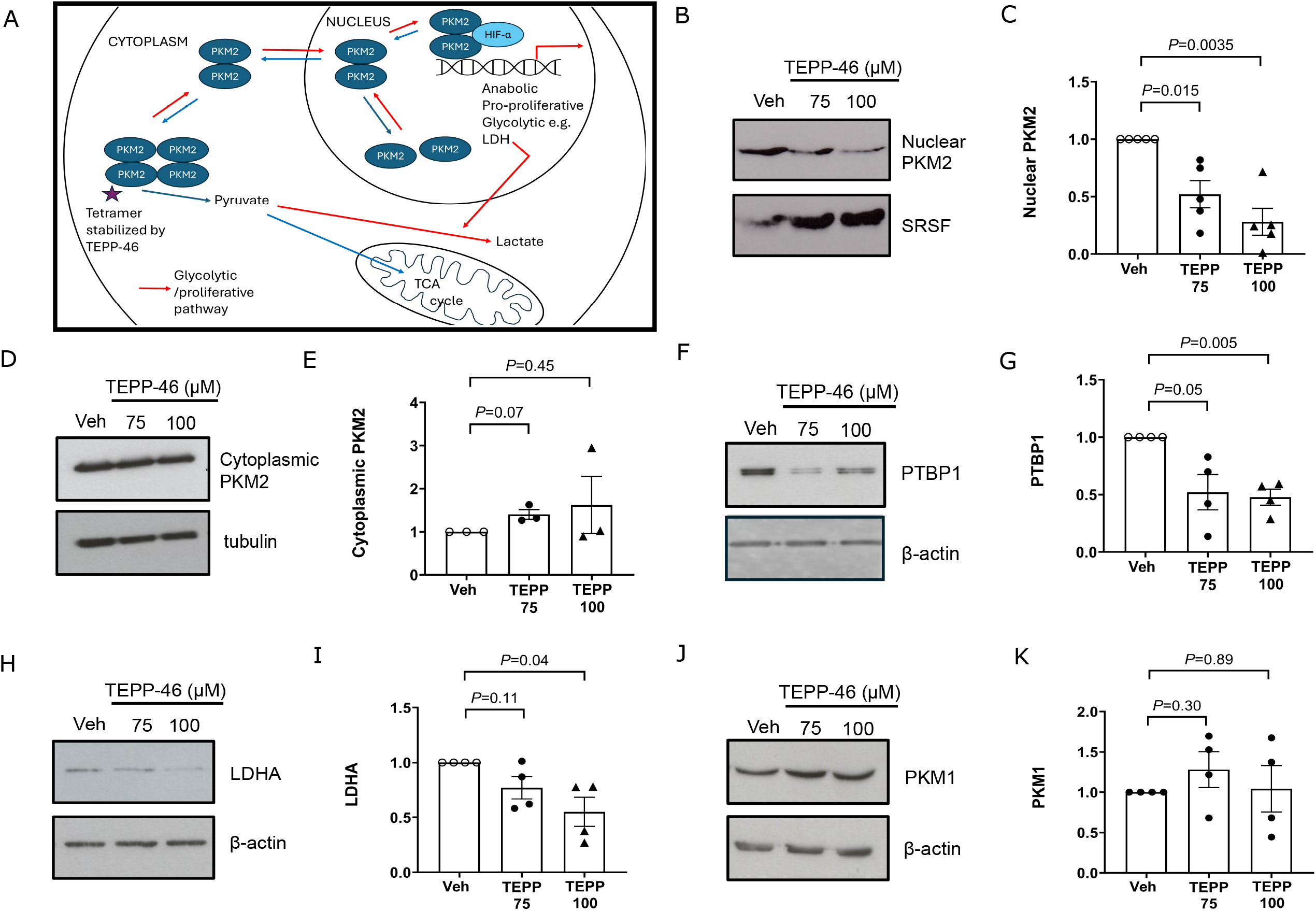
Interrogation of how TEPP-46 affects PKM2 levels and its pathways in BOECs. Human BOECs were isolated from the peripheral blood of healthy volunteers and grown as previously described(8), with approval from the local Research Ethics Review Committee (18/EE/0269). Panel A shows a schematic for the PKM2 axis. Panels (B-C) show a representative Western blot of the levels of PKM2 in the nucleus (B) and quantitation of densitometry (C). Panels (D-E) show a representative Western blot of the levels of PKM2 in the cytoplasm (D) and quantitation of densitometry (E). The PKM2 antibody used in (B) and (D) is the PKM2 (D78A4) rabbit monoclonal antibody (4053, Cell Signaling, USA) and the nuclear and cytoplasmic fractions were obtained using NE-PER^™^ nuclear and cytoplasmic extraction kit (78833, ThermoFisher Scientific, USA). Panels (F-I) show the levels of PTBP1 (anti-PTBP1 antibody [SH54], ab30317 from Abcam, UK); and LDHA (LDHA [C4B5] rabbit monoclonal antibody, 3582 from Cell Signaling, USA) in BOECs after 48 hours of treatment with either vehicle or TEPP-46. For these panels F and H show representative Western blots and panels G and I the corresponding densitometry. Panels (J-K) show the levels of PKM1 (PKM1 [D30G6] rabbit monoclonal antibody, 7067 from Cell Signaling, USA) in BOECs after 48 hours of treatment with either vehicle or TEPP-46 and quantitation of densitometry. In panels (C, E, G, I, and K) the mean is shown, and comparisons have been made using unpaired t-tests (for parametric data) or Mann-Whitney tests (for non-parametric data). TEPP-46 was the kind gift of Dr Craig Thomas. Antibodies used for loading controls include β-actin (A5441, Sigma-Aldrich, USA), tubulin (T5168, Sigma-Aldrich, USA), and SRSF1 (32-4600, Invitrogen, USA).

As there is evidence that TEPP-46 affects BOECs in a manner favoring oxidative phosphorylation, we decided to evaluate its effects in an animal model. The Sugen-hypoxia rat model was selected as it causes severe pulmonary hypertension and small artery occlusion mimicking human disease(9). The experimental protocol is summarized in Fig.2A. Oral treatment with TEPP-46 reduced the elevated right ventricular systolic pressures caused by exposure to Sugen-hypoxia (Fig.2B). TEPP-46 also ameliorated the body weight loss associated with Sugen-hypoxia (Fig.2C), and improved survival (Fig.2D). The decrease in the right ventricular mass indexed to body weight (Fig.2E) did not reach significance although interestingly, the increase in the left ventricular mass index was ameliorated by oral TEPP-46 (Fig.2F). There was no change in either cardiac output (Fig.2G) or ejection fraction (Fig.2H) between all groups.

**Figure 2:**
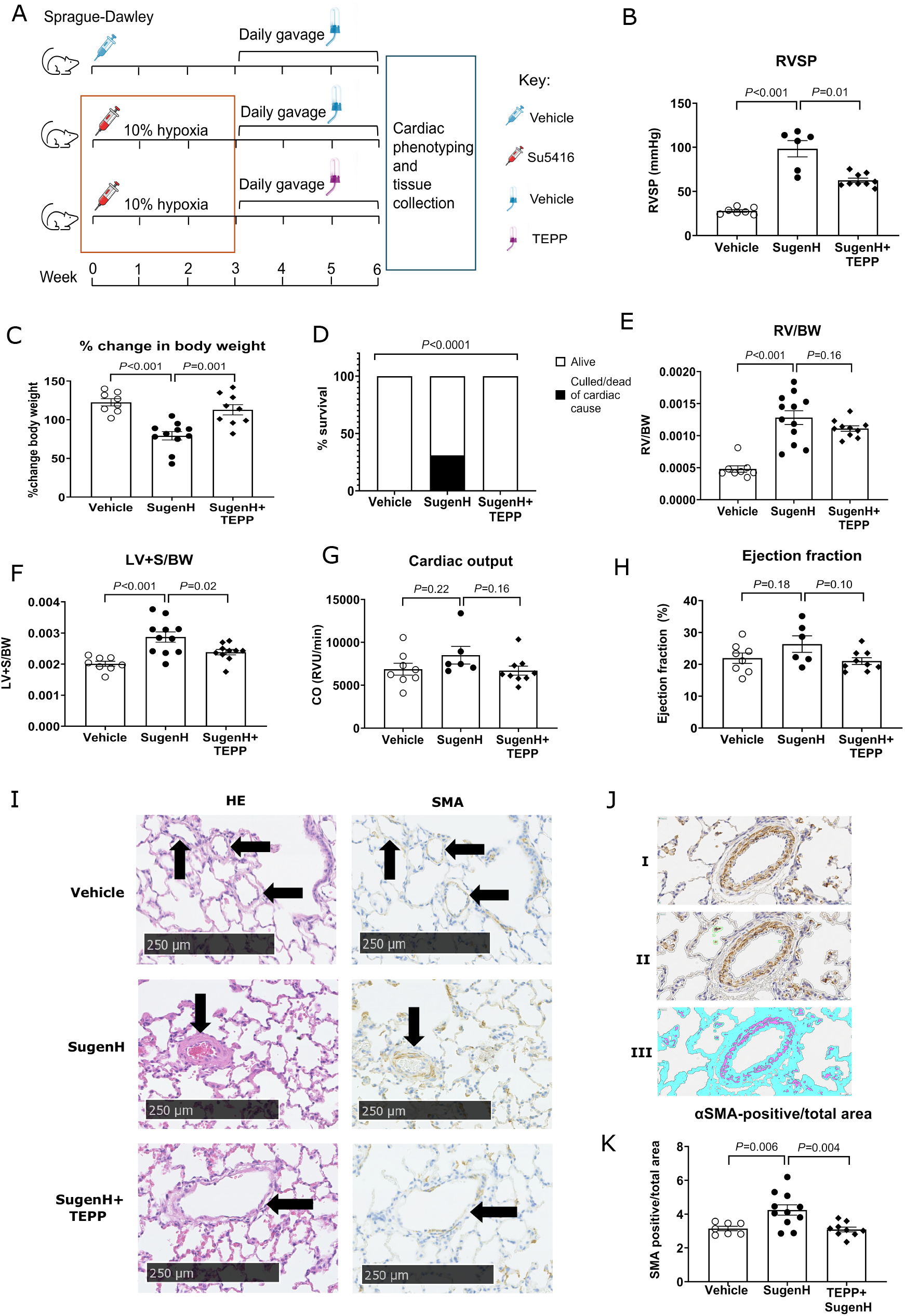
Characterization of TEPP-46 in a Sugen-hypoxia rat model. Panel A shows the experimental protocol, in which male Sprague-Dawley rats were exposed to either vehicle or 20mg/kg of Sugen-5416. Rats receiving Sugen-5416 were then exposed to hypoxia (10% oxygen) for 3 weeks. On exiting the hypoxic chamber, this cohort was split in two; with one group receiving daily gavage with 50mg/kg of TEPP-46 and the other receiving daily vehicle alone. Panels (B-D) show the right ventricular systolic pressures (RVSP, panel B), the change in body weight (panel C), and the survival of these animals as a percentage of their respective cohorts - defined as having to be culled due to reaching pre-determined severity protocols or death from a cardio-pulmonary cause (panel D). Panel E shows the right ventricular weight indexed to body weight (RV/BW). Panels (F-H) show the left ventricular weight indexed to body weight (LV/BW, panel F), the cardiac outputs (CO, panel G) and the ejection fraction (EF, panel H) of these rats. Panel I shows serial representative histological sections of the lungs of rats from each cohort, which have been stained for hematoxylin and eosin (HE) or α-smooth muscle actin (SMA). Black arrows show small blood vessels (≤250 micrometers). Panel J shows the Visiopharm algorithm to quantify the degree of SMA staining. Subpanels (I-II) show the initial histological section (I), and the delineation of tissue by the algorithm (dotted outlines, subpanel II). Subpanel (III) shows the identification of SMA positivity (pink), and of overall tissue area (turquoise). Panel K shows the result of that quantification in a graphical format. In panels (B-C, E-H and K) the mean is shown, and comparisons have been made using unpaired t-tests (for parametric data) or Mann-Whitney tests (for non-parametric data). In panel D the distribution of alive: culled or dead has been compared between groups using chi-squared testing. All animal procedures were performed following University of Cambridge guidelines, and in line with the Animals (Scientific Procedures) Act 1986 under the project license 70/8850.

Immunohistochemical examination revealed increased muscularization of the smaller vessels (50-200μm) in the lungs of the rats exposed to Sugen-hypoxia (Fig.2I) as shown by α-smooth muscle actin (SMA) staining. These changes were markedly improved in the rats exposed to Sugen-hypoxia which subsequently received TEPP-46. SMA-staining was quantified using Visiopharm software. In brief, the algorithm delineated all non-white pixels as ‘tissue’ and scored pixels above a threshold intensity for DAB-positivity as positive (Fig.2J). Therefore, an objective score can then be calculated for SMA-positivity per tissue area. Exposure to Sugen-hypoxia increased SMA-positivity and hence smooth muscle cell numbers, and this change was ameliorated by oral TEPP-46 (Fig.2K).

We show that pharmacological manipulation of the glycolytic process has beneficial effects in both a cell and rat model of pulmonary arterial hypertension. This is supported by evidence from previous studies. Increased PKM2 expression has been shown in the right ventricles of both rats subjected to pulmonary artery banding (PAB) and PAH patients. It is known that PARP1-dependent poly-ADP-ribose (PAR) is critical to the nuclear retention and function of PKM2. Shimauchi(10) showed that either genetic deletion of PARP1 or pharmacological blockade (olaparib, a PARP-1 inhibitor; or TEPP-46) was sufficient to improve markers of right ventricular function in rodents subjected to PAB.

Increased phosphorylated-PKM2 levels (which prevents tetramerization) and decreased tetrameric forms of PKM2 have been shown in both lung tissue and pulmonary artery smooth muscle cells derived from PAH patients(11, 12). In these pulmonary artery smooth muscle cells, TEPP-46 inhibits hypoxia-driven proliferation from both controls and PAH patients(12). Taken together with our study this suggests that the Warburg effect is significant in multiple cell types and organs in PAH models and patients. We conclude that antagonization of this effect would offer a novel therapeutic option for this deadly disease.

## Materials and methods

### Cell culture and reagents

Human BOECs were isolated from the peripheral blood of healthy volunteers and grown as previously described(8), with approval from the local Research Ethics Review Committee (18/EE/0269). TEPP-46 was the kind gift of Dr Craig Thomas. BOECs seeded as described in Table 1 were treated with vehicle + DMSO or TEPP-46 (75 or 100μM) in EGM2 (supplemented with 10% [v/v] FBS), for 48 hours, with fresh TEPP-46 treatment after 24 hours.

**Table 1:** BOEC seeding densities.

| Culture plate/dish | Cell numbers | Media volume per well/plate (ml) |
| --- | --- | --- |
| 6-well plate | 1.8-2x10<sup>5</sup> | 2 |
| 60 mm dish | 3.9-4x10<sup>5</sup> | 4.5 |

### Western blotting

BOECs cultured in 60 mm dishes were lysed with 200μl of radioimmunoprecipitation assay (RIPA) buffer supplemented with cOmplete^™^ protease inhibitor cocktail (Emerald Scientific, 1:19 dilution) on dry ice. The dishes were gently scraped and lysate collected into 1.5ml Eppendorfs. Samples were sonicated for 25-30 seconds and centrifuged (15,000 x G, 15 minutes, 4°C) before the clarified supernatant (protein lysate) was collected. For certain experiments, nuclear and cytoplasmic fractions were obtained using the NE-PER^™^ nuclear and cytoplasmic extraction kit (78833, ThermoFisher Scientific, USA). Protein concentration in the clarified lysate was quantified using a Bicinchoninic acid (BCA) Protein Assay according to manufacturer’s instructions (Bio-Rad). Protein concentration was determined using linear regression of a standard curve of bovine serum albumin (BSA). Samples were aliquoted into 20-80μg stocks and stored at −80°C until use.

Following this, samples were processed under reducing conditions, lysates were prepared for loading in laemelli buffer - 312.5 mM Tris-HCl, pH 6.8, 10% (w/v) sodium dodecyl sulphate, 50% (w/v) glycerol and 0.007% (w/v) bromophenol blue supplemented with 12.5% (v/v) β-mercaptoethanol to a 1:4 dilution; and heat-treated for 10 minutes at 95°C to achieve protein denaturation. Samples and PageRuler Plus Prestained Protein Marker (ThermoFisher Scientific) were loaded onto 12% polyacrylamide gels prepared in-house using National Diagnostics gel reagents or Novex NuPAGE^™^ 4-12% Tris-Glycine Gels (Invitrogen) in tanks containing SDS-running buffer. These gels were run at 100 V for 10 minutes, allowing samples to uniformly migrate through the lower density stacking gels into the higher density resolving gel, before running at 150 V for ∼1.5 hours or until the desired ladder separation was achieved. For proteins with a molecular weight up to 100kDa, gels were transferred to 0.2μM pore sized polyvinylidene difluoride (PVDF) membranes using the P0 setting on the iBlot® 2 Dry Blotting System (ThermoFisher Scientific) or onto membranes by manual semi-dry transfer (Amersham PVDF, 0.2μM pore size, GE Healthcare) for ∼1.5 hours, pre-activated by submersion in methanol. Proteins greater than 100kDa were transferred onto 0.2μM pore-sized PVDF membranes preactivated with methanol, via a wet-transfer system overnight (∼16 hours) at 4°C and 90mA.

After completion of transfer, membranes were blocked with 5% (w/v) skimmed milk in Tris-buffered saline (TBS; 6.05 g/l Tris-Base,8 g/l NaCl, pH 7.4) supplemented with 0.05% (v/v) Tween 20 (TBST) for 1 hour at room temperature under gentle rotation. Blots were washed with TBST 3 times and incubated with the primary antibodies listed in Table 2. Membranes were also incubated with appropriate loading controls for 1 hour at room temperature before washing and addition of secondary antibodies. Membranes were washed again with TBST before incubation with 5% milk (v/v) containing appropriate streptavidin-conjugated secondary antibodies (Dako) for 1 hour at room temperature (1:2000, Table 3).

**Table 2:**
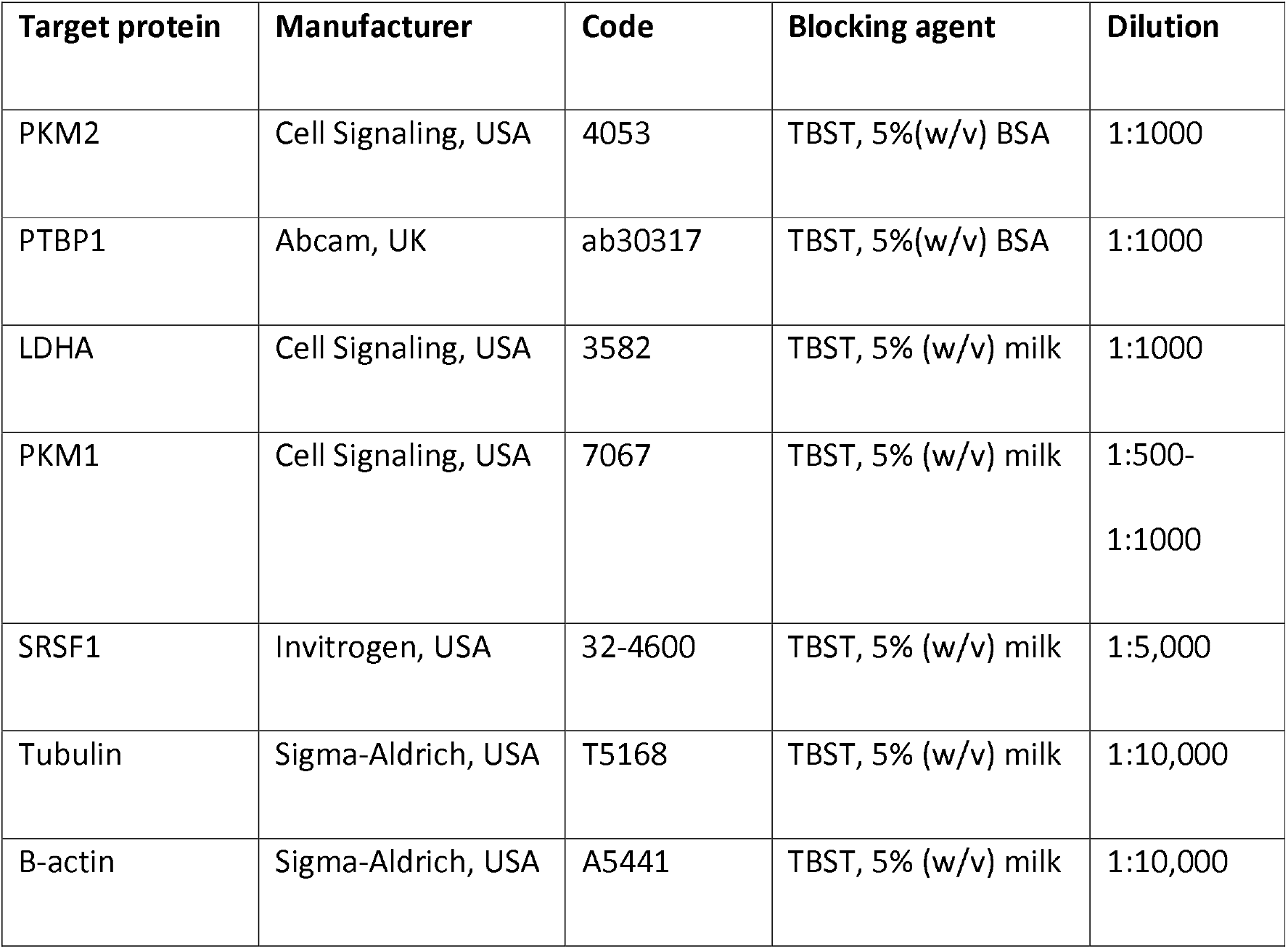
Primary antibodies used for Western blotting.

**Table 3:**
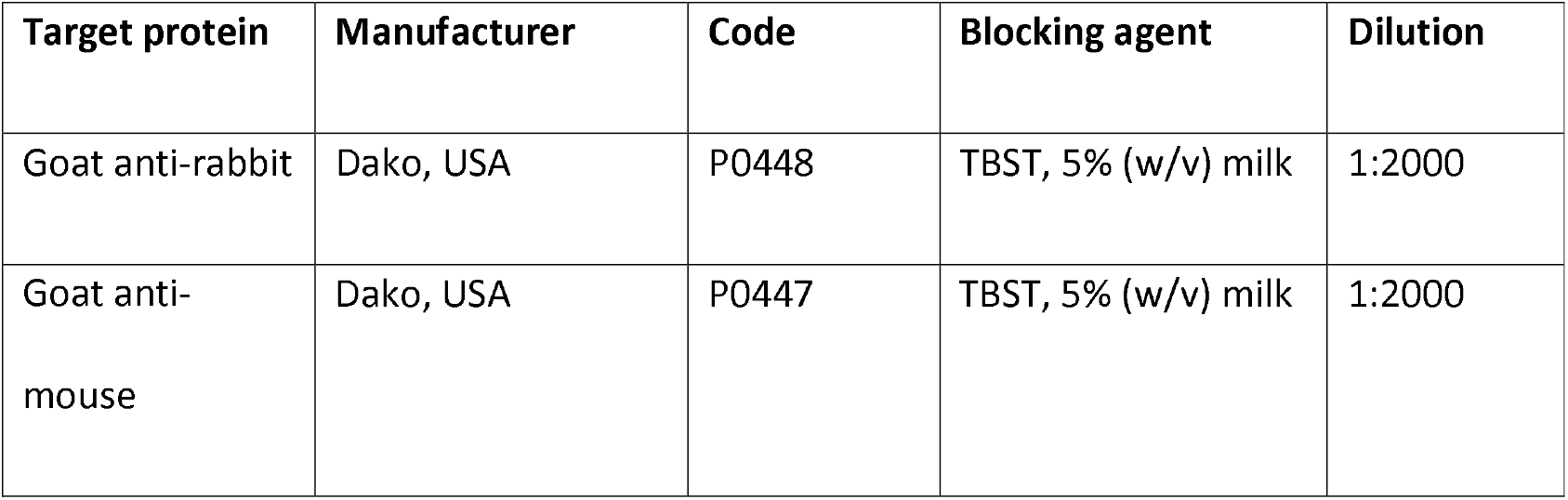
Primary antibodies used for Western blotting.

| Target protein | Manufacturer | Code | Blocking agent | Dilution |
| --- | --- | --- | --- | --- |
| Goat anti-rabbit | Dako, USA | P0448 | TBST, 5% (w/v) milk | 1:2000 |
| Goat anti-mouse | Dako, USA | P0447 | TBST, 5% (w/v) milk | 1:2000 |

### Rat Sugen-hypoxia model

All animal procedures were performed following University of Cambridge guidelines, and in line with the Animals (Scientific Procedures) Act 1986 under the project license 70/8850. Male Sprague Dawley rats (∼250-300g) were sourced from Charles River (UK) and allowed to acclimatize for 1 week. Power calculations were performed to determine the minimum group size required for sufficient statistical power (n=11, 20% effect change (1 sided) with 80% power and 95% confidence). Rats were randomly assigned to experimental groups (n=8-12 per group) and identified by tail number for experimental blinding purposes.

Rats were injected subcutaneously with VEGFR inhibitor Sugen-5416 (20mg/kg) or vehicle prior to entering hypoxia (10% oxygen) or normoxia for 21 days. Following hypoxic or normoxic exposure, rats were randomly allocated to oral gavage with either vehicle or 50 mg/kg TEPP 46 (5 mg/ml v/v in 10% DMSO v/v, 20% HP-β-CD g/v) daily for 21 days.

Approximately 24 hours following the final dosage, animals were subjected to right and left heart catheterization and tissue collection under terminal anesthesia. The right ventricle was dissected and weighed, as was the remaining left ventricle and septum. The right lung was snap-frozen in liquid nitrogen for RNA extraction and the left lung was isolated and perfused with 10% neutral buffered formalin (10% NBF) for 5 minutes, after which tissue was stored in 10% NBF before embedding into paraffin blocks.

